# The Interactions of Potassium Hydroxide in the Vicinity of Connective Disulfide Bonds of the Fab Regions of Some Human Immunoglubulins: Preliminary Computational Implications for Affinity on Cys-Cys Disulfide Bridges

**DOI:** 10.1101/2022.08.14.503886

**Authors:** Emin Zumrutdal

## Abstract

The average human lifespan continues to increase with the increase in data flow and the advancement of related technological developments. However, this development brings with it many diseases, including immunological problems. Immunoglobulin varieties found in different organisms in the last 3-4 decades continue to be hope for many diseases.

Interest has focused on the lesser weight but more mobile immunoglobulins found in camelids. Later, different types of these antibodies were tried to be made with biotechnological engineering and their effectiveness continues to be investigated. Disulfide bridges located on the immunoglobulin are one of the key points for the structure and function of the immunoglobulin. The interest of potassium hydroxide in disulfide bridges may enable us to damage or break these bonds. For this purpose, in this study, the relationship between disulfide bridges between light and heavy chains and potassium hydroxide was investigated. It was observed that the affinity of potassium hydroxide to disulfide bridges occurred exergonically. In the light of this information, it can be thought that lighter, more functional immunoglobulin fragments and nanobodies can be formed with potassium hydroxide compared to conventional immunoglobulin.

## INTRODUCTION

While the immunoglobulin(Ig) structure is preserved in mammals, it may differ between species. The Ig monomer consists of 2 heavy and 2 light chains(1). The disulfide bonds between the chains give Ig its 3D structure. These locations of disulfide bonds are nearly identical in all human Ig. In the respiratory system, IgA is usually found in the respiratory tract. However, it is known that other Ig such as IgG and IgM are present in addition to IgA(2–4)(Figure-1).

**Figüre-1:**
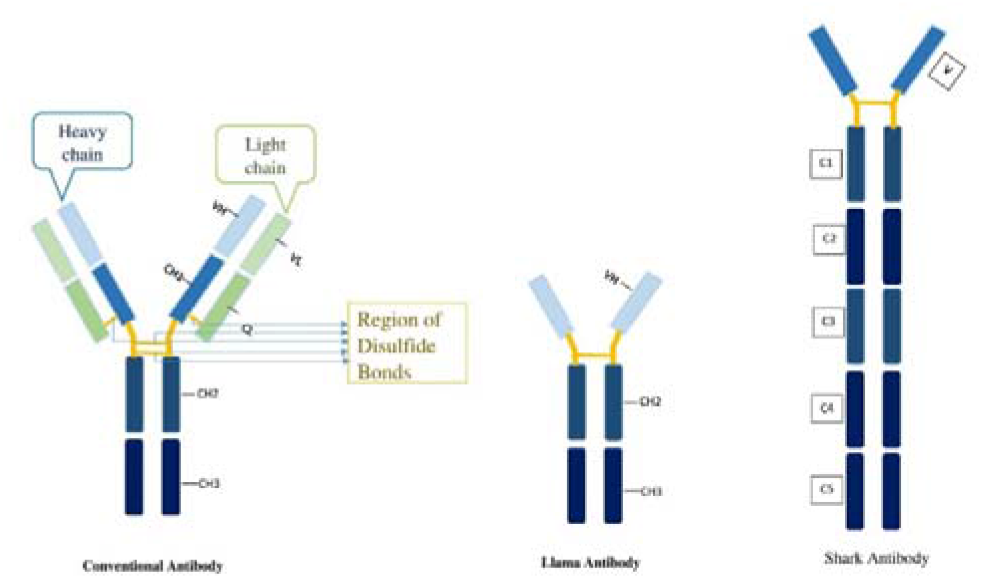
Different Ig structures in different species; human, llama, shark.

Ig mobility is crucial to its function. The rapid interaction of certain domains in the variable chain of Ig with the antigen enables the body to respond to diseases early. In light of this information, the activation of Ig fragments has received considerable attention(5).

Interest in Ig constructs has increased even more after the discovery of camelid heavy-chain antibodies(6,7). Different species can have different Ig structures(Figure-1). These antibody fragments, called by various names such as VHH fragment, single-domain antibody (sdAb) or nanobodies, can recognize the antigen. These structures have been used in many studies for therapeutic purposes. Today, usage areas are gaining more and more importance day by day(8–10).

The fact that Ig fragments formed by the breaking of disulfide bridges in Ig can display nanobody properties can create a different innovative dimension in studies.

In a previous study, it was observed that inhalation potassium hydroxide (KOH) administration positively supported oxidative stress in the lungs of mice(11). KOH is a reactive molecule that is interested in disulfide bonds(12).

In this study, we investigated the potential for formation of lighter Ig or nanobody than conventional Ig with this effect of KOH.

For this purpose, the affinity of KOH to the disulfide bonds between the heavy and light chains in IgA, IgG and IgM was investigated.

## Materials and Methods

### Ligand and Receptor retrieval

Immunoglobulin A (IgA), immunoglobulin G (IgG) and immunoglobulin M (IgM) 3D structures used as target receptors in this study were downloaded from the RCSB PDB database (https://www.rcsb.org/), along with codes 3QNX (IgA, resolution: 2.20 Å), 1AD9-1HZH (two different conformations of IgG with resolutions of 2.80 Å and 2.70 Å) and 1HEZ (IgM, resolution: 2.70 Å), respectively. In this docking study, potassium hydroxide (KOH) used as the ligand against human immunoglobulins was drawn using GausView 6.0.16 embedded in Gaussian 09 program and optimized with the B3LYP hybrid function (6-311G (d, p) basis set) (13).

In this molecular docking study performed to investigate the affinity of KOH with human immunoglobulin (Ig) molecules, the following hypothesis was established: Due to the reactive nature of the KOH molecule, it may cause the breakage of single disulfide bonds (-S—S-) bridging the heavy (H) and light (L) chains in the antigen-binding sites (Fab region) of human IgA, IgG and IgM antibodies. Indeed, the breakage of a covalent bond may not be demonstrated in docking simulations, however, the strength of the molecular affinity of KOH towards this region can be calculated. Thus, KOH, in solution, may pave the way for the light chain to be released and freely diffused in the human plasma. In line with this approach, prior to docking, Cys219-Cys133 (PDB ID: 3QNX), Cys127-Cys213/Cys230-Cys214 (PDB ID: 1AD9 and PDB ID: 1HZH, respectively) and Cys214-Cys135 (PDB ID: 1HEZ) residues in IgA, IgG and IgM molecules were visualized and targeted using Chimera 1.15 (Figure 2) (14). In addition to the single-ligand docking protocol, a simultaneous double-ligand docking protocol was also applied to observe the changes in the strength of the docking binding free energy (ΔG°=kcal/mol) in case the number of KOH molecules in the vacuum increases, and therefore, the affinity change of two KOH molecules in the vicinity of the disulfide bonds was also demonstrated. The simultaneous docking protocol is a fairly new approach in molecular docking discipline, thus, our study also provides new information to the literature in this respect.

**Figure 2.**
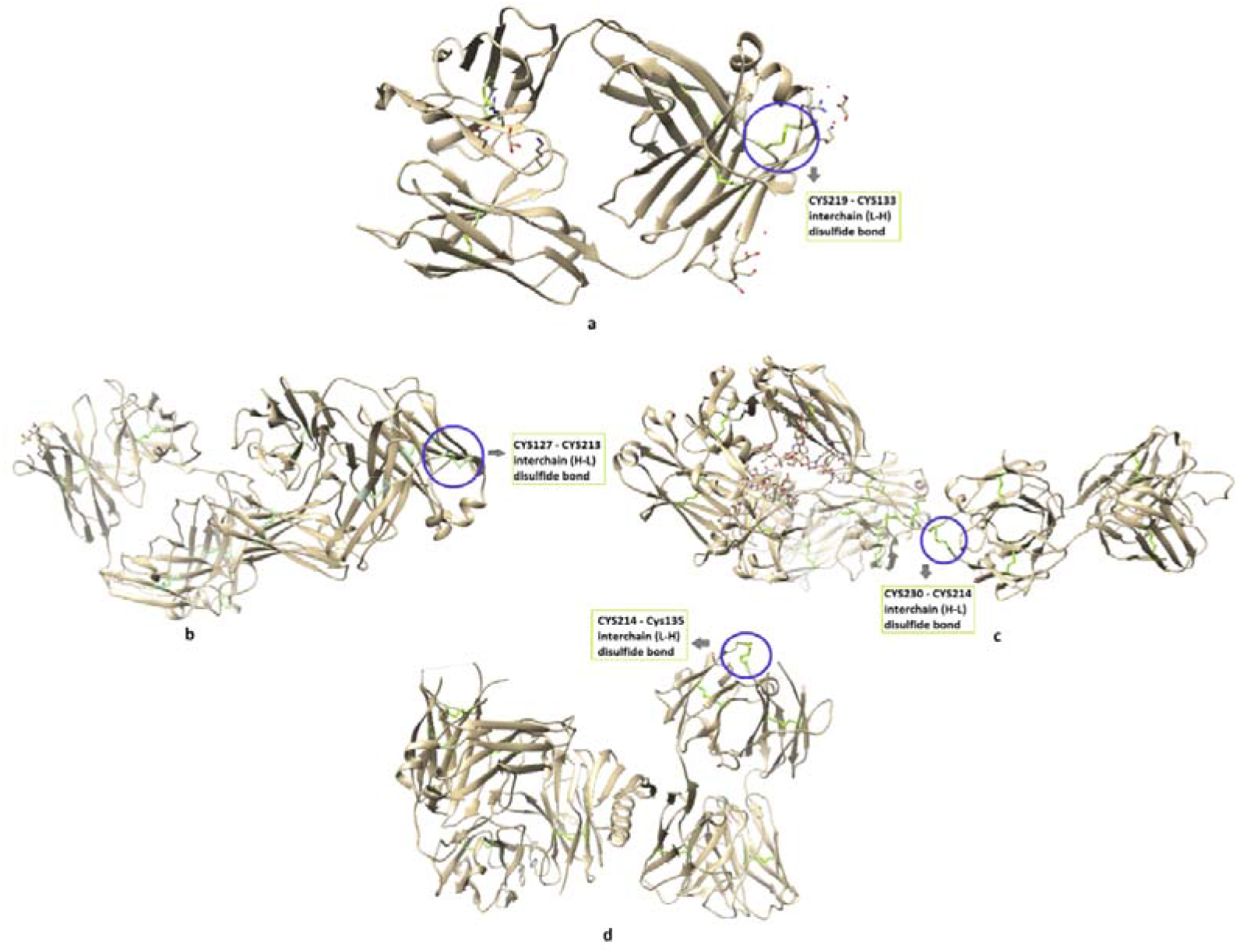
Cysteine residues (in blue circles) targeted at the Fab (antigen-binding site) sites of human immunoglobulin IgA (**a**), IgG (**b** and **c**) and IgM (**d**) molecules in molecular docking simulations and involved in the formation of disulfide bonds. Immunoglobulin proteins are depicted in the cartoon mode. Rendering was achieved using Chimera 1.15rc program.

### Molecular docking between potassium hydroxide and human immunoglobulin structures using single and simultaneous docking protocols

Prior to docking simulations with AutoDock Vina 1.2.0, target (IgA, IgG and IgM) and ligand (KOH) structures were prepared with AutoDockTools 1.5.6 and saved in pdbqt format (15,16). In molecular docking simulations with human Igs and KOH, polar hydrogen atoms in receptors and ligand molecules were retained, whereas, non-polar hydrogens were merged. Kollmann charges were assigned to the receptors and Gasteiger charges were assigned to the KOH. Rotatable bonds of KOH were set free during the docking calculations.

A grid box size of 5×5×5 Å points (x = 26.105; y = 10.749; z = 58.310) with 0.375 Å grid spacing was set for IgA, 4×4×4 Å points (x = 55.254; y = 82.653; z = −5.724) and 5×5×5 Å points (x = 73.242; y = 108.143; z = 157.489) with 0.375 Å grid spacing was set for each IgG conformations (1AD9 and 1HZH), and 5×5×5 Å points (x = 60.752; y = 74.466; z = 13.670) with 0.375 Å grid spacing was set for IgM in single docking protocols. These grid box sizes were set in a way that KOH molecule could easily interact with the amino acid residues in the immediate vicinity of the targeted interchain disulfide (-S—S-) bonds of each Ig molecule (Figure 2). In the simultaneous multiple ligand docking protocol, the same cartesian coordinates and the same grid box sizes were used against each Ig molecule.

After four separate docking runs against a total of four different Ig conformations (IgA, two IgGs and IgM, 20 dockings for each) of KOH, all potential binding modes of the ligand were clustered by AutoDock Vina 1.2.0 and were ranked based on the binding free energy (ΔG°; kcal/mol) of the ligand conformation which showed the lowest binding free energy against each Ig molecule. The top-scored docking conformations of the KOH calculated by AutoDock Vina 1.2.0 among different poses against the Ig receptors were rendered and analyzed using Discovery Studio Visualizer v16 (17).

## Results

### Human immunoglobulin docking studies of potassium hydroxide using new AutoDock Vina

We conducted molecular docking simulations to demonstrate the intermolecular interactions of potassium hydroxide (KOH) with the disulfide bonds bridging the heavy and light chains in the Fab domains (antigen-binding site) of the three human immunoglobulins (Ig) as well as the amino acid residues around these bonds. In addition, the molecular affinity (binding free energy [ΔG°]; kcal/mol) of KOH against these disulfide bond regions targeted on Fabs of Igs was also calculated.

Considering the single KOH docking results, it was determined that KOH formed a hydrogen (H) bond with Leu132, metal-acceptor interaction with Cys133, very close to the disulfide bond (Cys219-Cys133) in the Fab region of human IgA, and the calculated binding free energy (ΔG°) of KOH against this targeted region is −1.55 kcal/mol, which is energetically favorable (Table 1, Figure 3A).

**Table 1.** Predicted docking binding free energies and human immunoglobulin amino acid-interaction results for potassium hydroxide (KOH).

| Compound | Receptor | $\Delta G_{\text{best}}$<br>(kcal/mol) | Classical<br>H-bond | Non-<br>classical<br>H-bond | Electrostatic | Other<br>(metal-acceptor) |
| --- | --- | --- | --- | --- | --- | --- |
| *Single KOH docking* |  |  |  |  |  |  |
| KOH | IgA | -1.55 | Leu132 | - | - | Cys133 |
| KOH | IgG<br>(1AD9) | -0.85 | Asn209 | - | Glu212,<br>Cys213 | - |
| KOH | IgG<br>(1HZH) | -1.11 | Lys228 | - | Cys214 | Cys230 |
| KOH | IgM | -1.36 | Asn137 | - | Glu213 | Asn210, Cys214 |
| *Multiple KOH docking* |  |  |  |  |  |  |
| KOH+KOH | IgA | -3.06 | - | - | - | Cys133 |
| KOH+KOH | IgG<br>(1AD9) | -2.67 | Glu212 | Ser128 | Glu212,<br>Cys213 | Cys213 |
| KOH+KOH | IgG<br>(1HZH) | -1.81 | Cys214 | - | Cys214 | Lys228, Cys230 |
| KOH+KOH | IgM | -2.66 | Asn137 | - | Glu213,<br>Cys214 | Asn210, Cys214 |

IgA: Human immunoglobulin A Fab domain (PDB ID: 3QNX) IgG: Human immunoglobulin G Fab domain (PDB ID: 1AD9 and 1HZH) IgM: Human immunoglobulin M Fab domain (PDB ID: 1HEZ) $\Delta G_{\text{best}}$ : most favorable binding free energy (kcal/mol).

**Figure 3.**
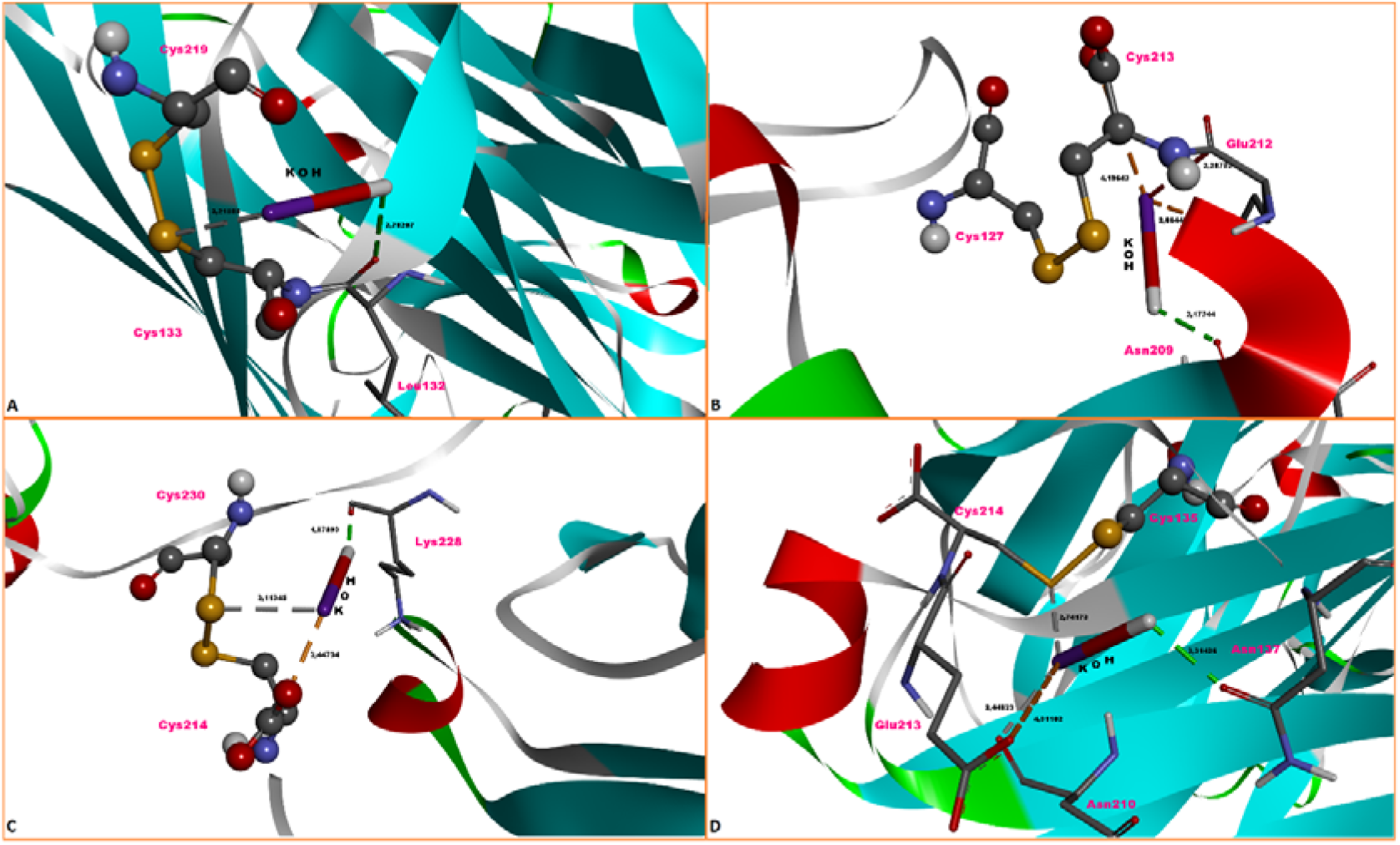
Single-ligand docking top-ranked conformations of potassium hydroxide (KOH) in complex with the Fab domains of human immunoglobulin A, G (two conformations) and M. **A**. IgA—KOH complex, **B**. IgG—KOH complex (PDB ID: 1AD9), **C**. IgG—KOH complex (PDB ID: 1HZH), **D**. IgM—KOH complex. The green and light cyan dashed lines represent hydrogen bonds, grey dashed lines represent metal-acceptor interactions, and the orange dashed lines represent electrostatic interactions.

The interactions of KOH in the vicinity of the targeted disulfide bond (Cys127-Cys213) region of human IgG Fab fragment consist of an H-bond with Asn209, and an electrostatic attraction interaction with Glu212 and Cys213. The calculated binding affinity of KOH for the IgG Fab fragment, although not highly strong, is energetically favorable (−0.85 kcal/mol) (Table 1, Figure 3B).

Amino acid interactions of KOH around the disulfide bond (Cys230-Cys214) that connects the constant heavy (C_H_) and constant light (C_L_) chains (in the Fab region) of IgG B12 (PDB ID: 1HZH), the other IgG molecule in our study, are given in Table 1 and Figure 3C. KOH formed a H-bond with Lys228, an electrostatic interaction with Cys214 and a metal-acceptor interaction with Cys230 in the vicinity of the Cys230-Cys214 disulfide bond (Figure 3C). The interaction strength of KOH around this disulfide bond is favorable and was calculated as −1.11 kcal/mol (Table 1). When the interaction of KOH around the disulfide bond (Cys214-Cys135) in the Fab region of the human IgM molecule is examined, it appears to have formed a H-bond with Asn137, an attractive charge interaction with Glu213, and a metal-acceptor interaction with both Asn210 and Cys214 (Table 1, Figure 3D). The calculated binding affinity of KOH in this region of the Fab domain is favorable (−1.36 kcal/mol) (Table 1). When the interactions of KOH with the relevant disulfide bonds in the Fab regions of IgA, IgG and IgM molecules in single docking simulations are examined (Table 1), it is concluded that KOH forms molecular contacts with at least one of the Cys residues in all Cys-Cys disulfide bonds which are targeted in docking. In addition, based on the binding free energy of the complexes, it was determined that disulfide bonds with the highest affinity for interaction with KOH (hence may be more open to physical perturbation) were detected in IgA and IgM (Table 1).

In Table 1, the interactions of KOH (in multiple form) with the targeted Cys-Cys disulfide bridges in the Fab domains of human immunoglobulins as well as its interactions with the amino acid residues in close proximity to these bonds, along with the binding free energy values are given. KOH molecules, in multiple docking, have formed two metal-acceptor interactions with Cys133 of IgA and the calculated free energy (ΔG°) of this binding is −3.06 kcal/mol (Table 1, Figure 4A). Compared to single KOH docking (Table 1), the combined binding energy of KOHs in multiple form around the targeted Cys219-Cys133 disulfide bond was dramatically increased. This finding shows that KOH molecules, more than one, can react energetically more efficiently in this region of IgA (Figure 4A).

**Figure 4.**
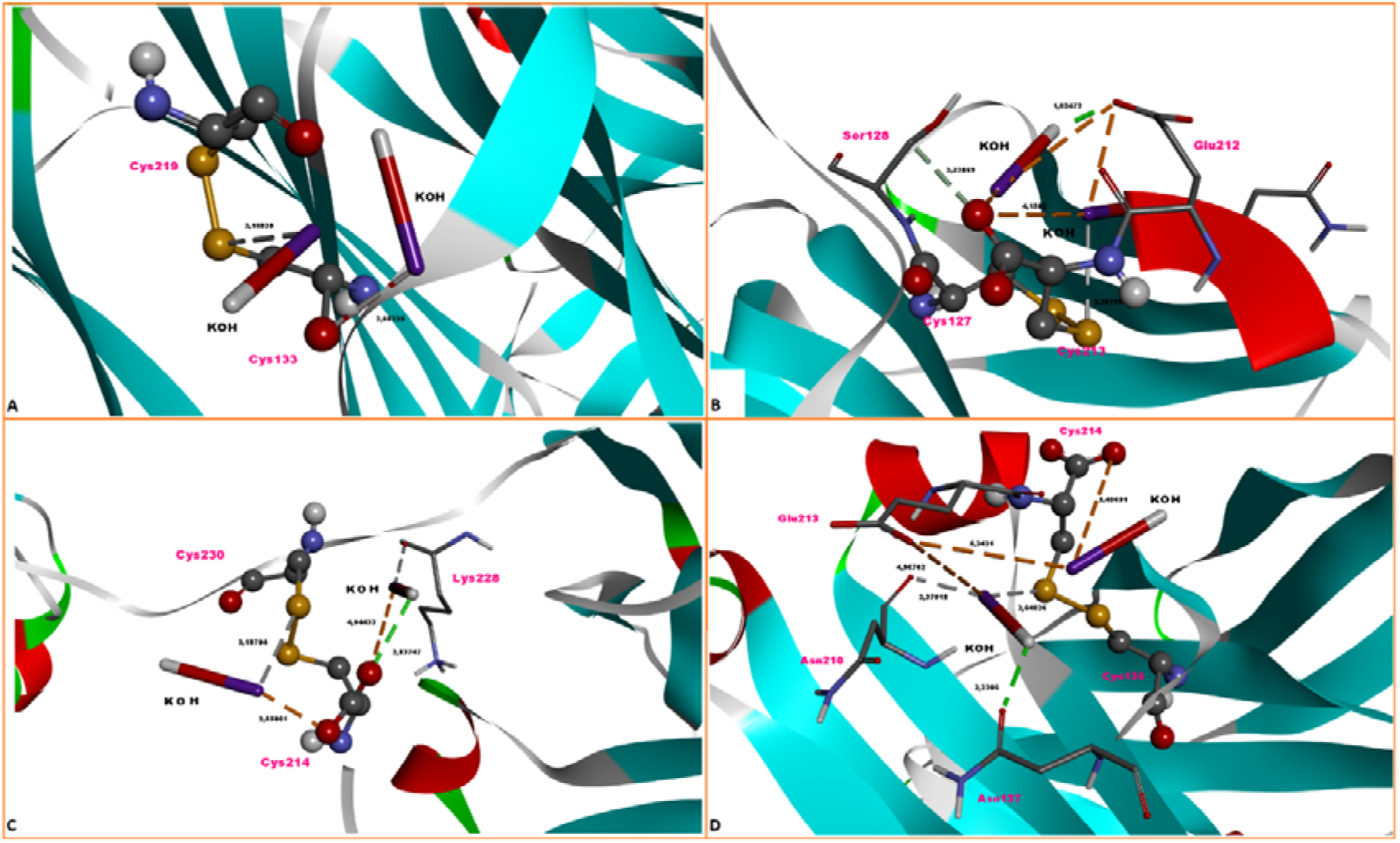
Multiple-ligand docking top-ranked conformations of KOH in complex with the Fab domains of human immunoglobulin A, G (two conformations) and M. **A**. IgA—KOH complex, **B**. IgG—KOH complex (PDB ID: 1AD9), **C**. IgG—KOH complex (PDB ID: 1HZH), **D**. IgM—KOH complex. The green and light cyan dashed lines represent hydrogen bonds, grey dashed lines represent metal-acceptor interactions, and the orange dashed lines represent electrostatic interactions.

KOHs formed two H-bonds, two attractive electrostatic interactions and a metal-acceptor interaction with Glu212, Ser128 and Cys213 residues of IgG in multiple docking, respectively (Table 1, Figure 4B). The calculated combined binding energy of these interactions is −2.67 kcal/mol, and increased dramatically (more than 2 fold) compared to single ligand docking (Table 1). In this multiple docking, the interactions of KOH molecules are too close to the targeted disulfide bond(Cys127-Cys213) and the sulfur atom of Cys213 is involved in this interaction(Figure 4B).

As the docking simulation of two KOH molecules against the other conformation (PDB ID: 1HZH) of the human IgG protein is evaluated, it is evident that KOH forms a H-bond, two attractive electrostatic and two metal-acceptor interactions with Cys214, Lys228 and Cys230 residues, respectively(Table 1, Figure 4C). Compared with single docking (Table 1), the combined binding strength (−1.81 kcal/mol) of the two KOH molecules at the targeted IgG site (Cys230-Cys214) was found to be slightly increased. In this docking, the KOH molecules are located very close to the target disulfide bond, and one of the KOHs formed a metal-acceptor interaction with the sulfur atom of Cys230 (Figure 4C).

In the targeted disulfide bond region of IgM (Cys214-Cys135), KOH molecules formed a H-bond, three attractive electrostatic interactions, and two metal-acceptor interactions with residues Asn137, Glu213, Cys214, and Asn210, respectively (Table 1, Figure 4D). One of the metal-acceptor interactions was occured between KOH and the sulfur atom of Cys214 (Figure 4D). Combined binding energies of KOH molecules at the site of the targeted disulfide bond are highly favorable (−2.66 kcal/mol), which is approximately twice as high as for single KOH docking (Table 1). When the results of multiple KOH docking are evaluated, it can be concluded that as the number of KOH molecules around the targeted disulfide bonds in Fab regions increases, the interaction of KOH with amino acids and disulfide bonds in these regions becomes stronger in a highly favorable manner (Table 1). This indicates that the collision effect of multiple KOHs on disulfide bonds (and possibly bond breakage) may greatly increase under biological conditions.

## DISCUSSION

Ig is a molecule in glycoprotein structure. In the 3D folded structures of Igs, the disulfide bonds are usually in contact with the outer surface. Therefore, it is suitable for external influences. Disulfide bonds are more specific between CH1 and CH2, between CH2 and CH2, and between CH1 and CL(Figure-1).

About 30 years ago, Igs lacking the light chain found in camelids, but functioning, broke new horizons by breaking down all stereotypes. Camelid antibodies carry heavy chain only. A different type of these molecular structures are also found in sharks(Figure-1)(18–20).

The absence of CH1 and light chain structures in these antibody constructs makes these antibodies lighter and more tissue penetration(21).

With the developing technology, these molecular structures consisting of only the VHH group are called single-domain antibodies (sdAbs) or nanobody. The longer CDR3, which appears with the nanobody, can more easily reach the antigen epitopes that classical antibodies cannot reach(22).

There is very little information about nanobodies yet. However, the efficiency of the structures formed by the breaking of Ig disulfide bonds as nanobody is worth investigating. Breaking of disulfide bonds can increase chain mobilization within the light and heavy chain as well as breaking down the Ig structure. This increases its ability to bind to epitopes.

The breaking of disulfide bonds begins with the affinity of different molecular structures such as KOH to these bonds. This study focused on IgA, IgG and IgM types due to their incidence.

KOH is a chemical that has been used as a caustic agent for years. Misuse in humans has led to fatal consequences(23,24). Hypochlorous acid (HOCl) is used as a cleaning material in almost every person’s home. However, HOCl has the same dangerous, even deadly effect. But would it be possible to survive without HOCl? HOCl is very important in the body defense system(25,26). HOCl is a chemical molecular agent that is produced in humans every hour, every day. HOCl used as a cleaning material in homes and HOCl used by leukocytes in defense have the same molecular structures. The difference is only in concentration. Some of the defense cells in the human body are like a constantly working factory for HOCl.

Could the same be true for KOH? Can we turn this molecular agent, which people have been afraid of for years, to the benefit of humanity by using it in harmony with tissue?

Molluscum Contagiosum was treated with KOH contact application to the skin(27,28). However, there is no inhaler use of KOH in humans. Very remarkable results have been obtained in studies on the antiviral effectiveness of KOH. In this study, KOH was buffered at the highest pH (0.9% NaCl solution, pH range 8.9-9.1) that can be taken orally and inhaled(11). This solution is compatible with interstitial tissue fluid.

In the study with KOH, KOH was administered as an inhaler and an oral spray. In vitro studies have shown the mucolytic activity of KOH in respiratory human mucus samples with its alkalizing, surface tension reducing, mucus contact angle reducing effect. In addition, exergonic binding energies to the envelope glycerophosphate structure for enveloped viruses, to the Coronavirus spike glycoprotein active site, to the human AGE2 active site and to the neurominidase active site for the Hemophilus Influenza virus were determined(11).

It has been shown in in vivo studies that studies with KOH prolong the life span in human fibroblast cell culture at different stages; It has been determined that it does not cause histopathological findings in oral tissue and lung tissue in young and old mice, positively affects oxidative stress markers and has a mucolytic effect in bronchoalveolar lavage(11).

In a previous KOH study, KOH inhalation decreased malondialdehyde levels and increased superoxide dismutase levels in aged mice, and the absence of histological pathological findings with KOH administration may support the functional part of the proteomic study(11).

Proteomic studies of therapeutics used in the treatment of diseases are important for elucidating the functions of molecules.

In this study, the interaction between the molecule (KOH) and the substrate (Ig,Disulfide bonds) was revealed. This work forms part of the structural proteomics study(Table-1).

In this study, disulfide bonds between KOH, CH1H and CL interacted exergonically in IgA, IgM and IgG(Figure-2). As the number of KOH molecules increases in the medium, this interaction increases even more(Table-1). This affinity suggests that KOH has a potential effect on breaking disulfide bonds. This result confirms our hypothesis.

**Figure-3:**
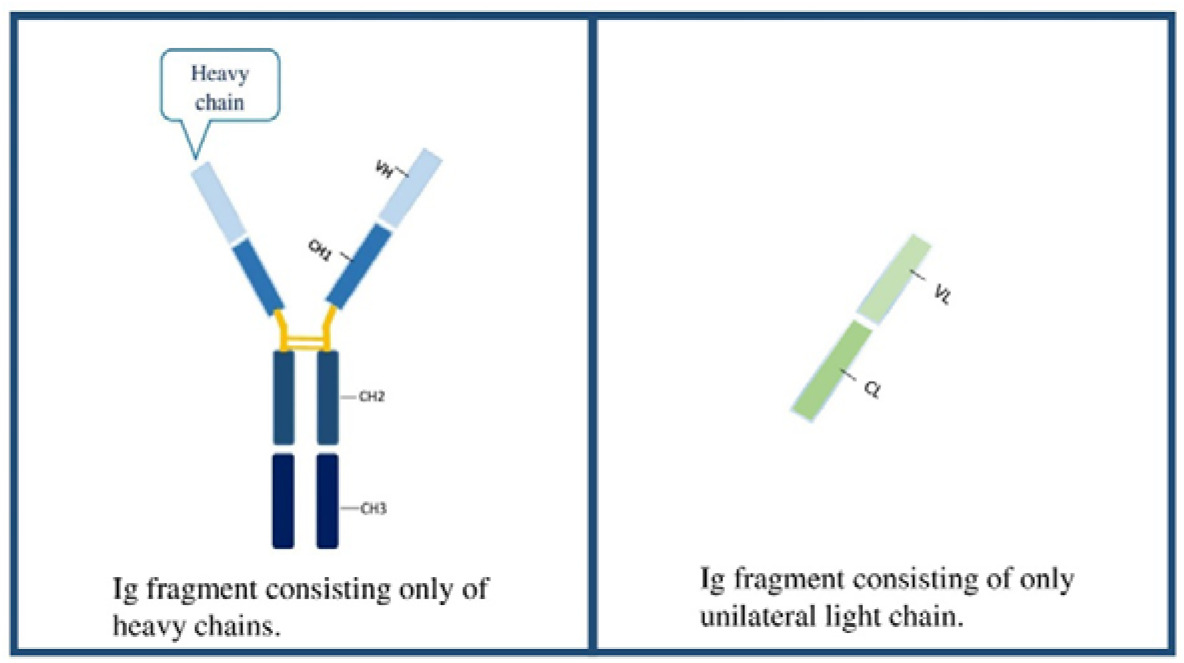
Potential light-weight Ig fragments that can be generated from conventional Ig by inhalation of KOH in the lungs.

Research with KOH should be supported by comparative (expression) proteomics, molecular dynamics and functional proteomics studies.

The interaction of KOH with Ig can lead to the formation of new nanobodies and the lung may be the nanobody building center. Nanobodies consisting of all Ig in the secretion can spread throughout the body by capillary circulation. If the KOH-disulfide bond interactions are strong enough to break the bond, our chance to see nanobodies of different sizes and functions increases.

Considering the 3-dimensional structure of Ig in the potential nanobady formation with KOH, the products that can be formed can push the limits of our imagination. The angulations in the Y region can also change with the effect of the disulfide bond in the CH1H, VH, CL and VL chain structures.

## CONCLUSION

In this study, the affinity of KOH to the disulfide bonds between the heavy and light chains in the Fab fragment of conventional immunoglobulin A,M and G has been demonstrated exergonically. It was determined that the Gibbs energies increased from −1 kcal/mol to −3 kcal/mol as the number of KOH molecules increased in the medium.

If nanobody formation from Ig or changes in 3D structures can be achieved with KOH inhalation, many immunological disease-causing epitopes can be detected very early.

KOH inhalation can be an effective treatment model for the people of the world with its easy production technology, low cost and being feasible in almost every country in the world.

